# Integrating global patterns and drivers of tree diversity across a continuum of spatial grains

**DOI:** 10.1101/367961

**Authors:** Petr Keil, Jonathan M. Chase

**Affiliations:** German Centre for Integrative Biodiversity Research (iDiv), Halle-Jena-Leipzig, Deutscher Platz 5e, 04103 Leipzig, Germany; Institute of Computer Science, Martin-Luther University Halle-Wittenberg, 06099 Halle (Saale), Germany

## Abstract

What drives biodiversity and where are the most biodiverse places on Earth? The answer critically depends on spatial scale (grain), and is obscured by lack of data and mismatches in their grain. We resolve this with cross-scale models integrating global data on tree species richness (S) from 1338 local forest surveys and 287 regional checklists, enabling estimation of drivers and patterns of biodiversity at any desired grain. We uncover grain-dependent effects of both environment and biogeographic regions on S, with a positive regional effect of Southeast Asia at coarse grain that disappears at fine grains. We show that, globally, biodiversity cannot be attributed to purely environmental or regional drivers, since regions are environmentally distinct. Finally, we predict global maps of biodiversity at two grains, identifying areas of exceptional species turnover in China, East Africa, and North America. Our cross-scale approach unifies disparate results from previous studies regarding environmental versus biogeographic predictors of biodiversity, and enables efficient integration of heterogeneous data.

---

What drives global variation in the numbers of species that live from place to place? For example, why are there fewer than 100 species of trees that live in millions of km^2^ in the boreal forests of Eurasia or North America^1^, while there can be hundreds of species co-occurring in as little as 50 ha in tropical forests of South America and Asia^2^?

The most important obstacle to answering these fundamental questions is a lack of data, especially in places with the highest biodiversity^3,4^. But even in the regions and taxa which have been well-sampled, the data are a heterogeneous mixture of point observations, survey plots, and regional checklists, all with varying area and sampling protocol^4^. For example, for trees, there are hundreds of 0.1 ha Gentry forest plots mostly in the New World^5^, hundreds of 1 ha ForestPlots.net plots throughout tropical forests^6^, dozens of > 2 ha CTFS-ForestGEO plots (www.forestgeo.si.edu), hundreds of published regional checklists^7^, and hundreds to thousands of other published surveys and checklists scattered throughout the published and grey literature. These together hold key information on global distribution of tree biodiversity, yet the lack of methods to address differences in sampling have so far prevented their integration.

Further, as could be said for many problems in ecology, attempts to map global biodiversity and to assess its potential drivers are severely complicated by the issues of spatial scale^8–11^: The most straightforward issue is the non-linear increase of number of species (S) with area^12^, which is why patterns of biodiversity cannot be readily inferred from sampling locations of varying area. The second issue concerns sets of sampling locations that do have a constant area (hereafter grain); even then a spatial pattern of S observed at a small grain may differ from a pattern at large grain^13–15^ – an example is grain-dependence of latitudinal diversity gradient^16^ [but see ref^17^]. The reason is that beta diversity (the ratio between fine-grain alpha diversity and coarse-grain gamma diversity) varies over large geographic extents^18^. Finally drivers and predictors of diversity have different associations with S at different grains^19–22^. For example, at global and continental extents, the association of S with topography increases with grain in Neotropical birds^22^ and the association with temperature increases with grain in global vertebrates^21^ and eastern Asian and North American trees^23^. Thus, biodiversity should ideally be studied, mapped, and explained at multiple grains^14^.

Although the abovementioned scaling issues are well-known^13,19,24,25^, methods are lacking that explicitly incorporate grain-dependence within a single model, allowing cross-grain inference and predictions. Furthermore, it is still common to report patterns and drivers of biodiversity at a single grain, resulting in pronounced mismatches of spatial grain among studies, and hindering synthesis. An example is the debate over whether biodiversity is more associated with regional proxy variables for macroevolutionary diversification and historical dispersal limitation, or with ecological drivers that include climatic and other environmental drivers, as well as biotic interactions^25–29^. While climate and other ecological factors usually play a strong role [but see ref^30^], studies differ in whether they view residual regional forces being weak^31–33^ or strong^34–36^. Even within the same group of organisms – trees – there is debate regarding whether environment^23,37–40^ or regional history^41–44^ drive global patterns. And yet, these studies are rarely done at a comparable spatial grain, and perhaps not surprisingly, studies from smaller plot-scale analyses^39,40^ typically conclude a strong role for environmental variation, whereas large-grain analyses^43,45^ show a strong role of historical biogeographic processes.

Here, we propose a cross-grain approach that allows estimation of contemporary environmental and regional predictors, as well as global patterns, of tree species richness across a continuum of grains, from plots of 10 × 10 m^2^ up to the entire continents. Our study has three main goals: (i) by explicitly considering spatial grain as a modifier of the influence of ecology versus regional biogeography, we aim to synthesize results among studies, and illustrate how the importance of these processes varies with grain. Apart from the well-known grain-dependent effects of environment, we also focus on the so far overlooked grain-dependent effects of biogeographic regions. (ii) The novelty of the approach is to model grain-dependence of every predictor (spatial, regional, or ecological) within a single model as having a statistical interaction with area, which enables integration of an unprecedented volume of heterogeneous data from local surveys and country-wide checklists – although such interaction has been occasionally tested^16,17,36^, to our knowledge it has not been applied to both spatial and environmental effects, nor for data integration and cross-grain predictions. (iii) We take the advantage of being able to predict biodiversity patterns at any arbitrarily chosen grain and we map the estimates of alpha, beta, and gamma diversity of trees across the entire planet.

## Results and Discussion

### Macroecological patterns

To explain the observed global variation of tree diversity (Fig. 1), we specified two models that predict S by grain-dependent effects of environmental variables (Supplementary Table 1), but differ in the way they model the grain-dependent regional component of biodiversity: model REALM attributes residual variation of S to location’s membership within a pre-defined biogeographic realm [as in ref^46^], while model SMOOTH estimates the regional imprints in S directly from the data using smooth autocorrelated surfaces. Both models explain more than 90% of deviance of the data (Supplementary Table 2) and both predict S that matches the observed S (Supplementary Fig. 1). This is in line with other studies from large geographical extents, where 70–90% model fits are common even for relatively simple climate-based models^23,40,46–48^.

**Figure 1.**
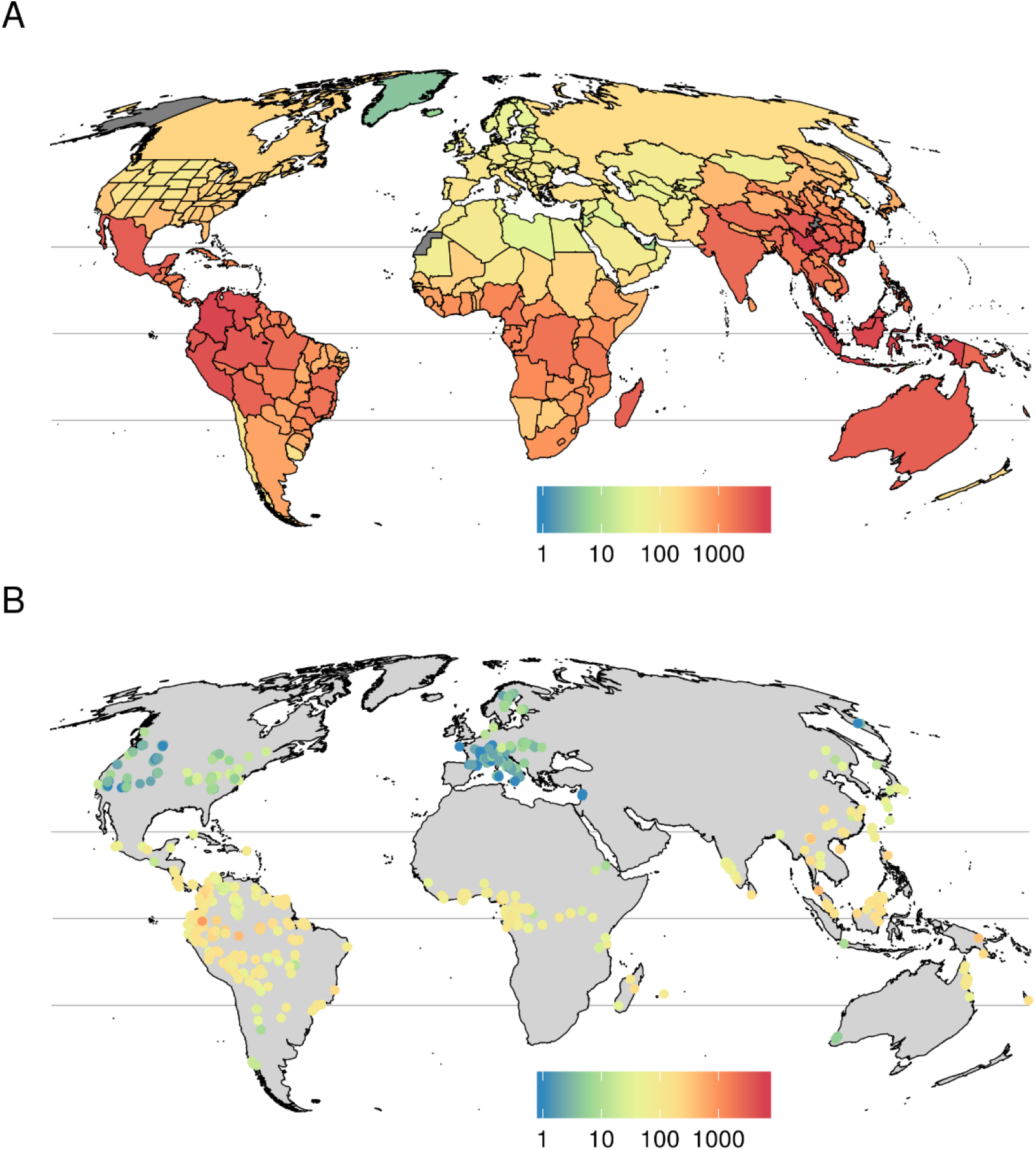
Raw data on observed tree species richness S (log_10_ scale) (A) at the country/states grain with 287 spatial units, and (B) at the plot grain with 1338 plots. Maps use Mollweide projection.

Next, we used model SMOOTH to predict patterns of S and beta diversity over the entire mainland, at a regular grid of large hexagons of 209,903 km^2^ and at a grid of local plots of 1 ha (Fig. 2A-C). We predict latitudinal gradient of S at both grains (Fig. 2A, B, Supplementary Fig. 2), which matches the traditional narrative in trees^49^ and other groups [ref^50^, p. 662–667]. However there are also differences between the patterns at the two grains, particularly in China, East Africa, and southern North America (Fig. 2C), where the plot-grain S is disproportionally lower than what would be expected from the coarse-grain S. These are regions with exceptionally high beta diversity and are in the dry tropics and sub-tropics with high topographic heterogeneity – examples are Ethiopian Highlands and Mexican Sierra Madre ranges, which have sharp environmental gradients and patchy forests, resulting in relatively low local alpha diversity but high regional gamma diversity. The exception is the predicted high beta diversity in China, where the historical component of beta diversity dominates the effect of environmental gradients (Fig. 2C vs F), as also suggested by refs^23,41,51,52^, and as discussed below.

**Figure 2.**
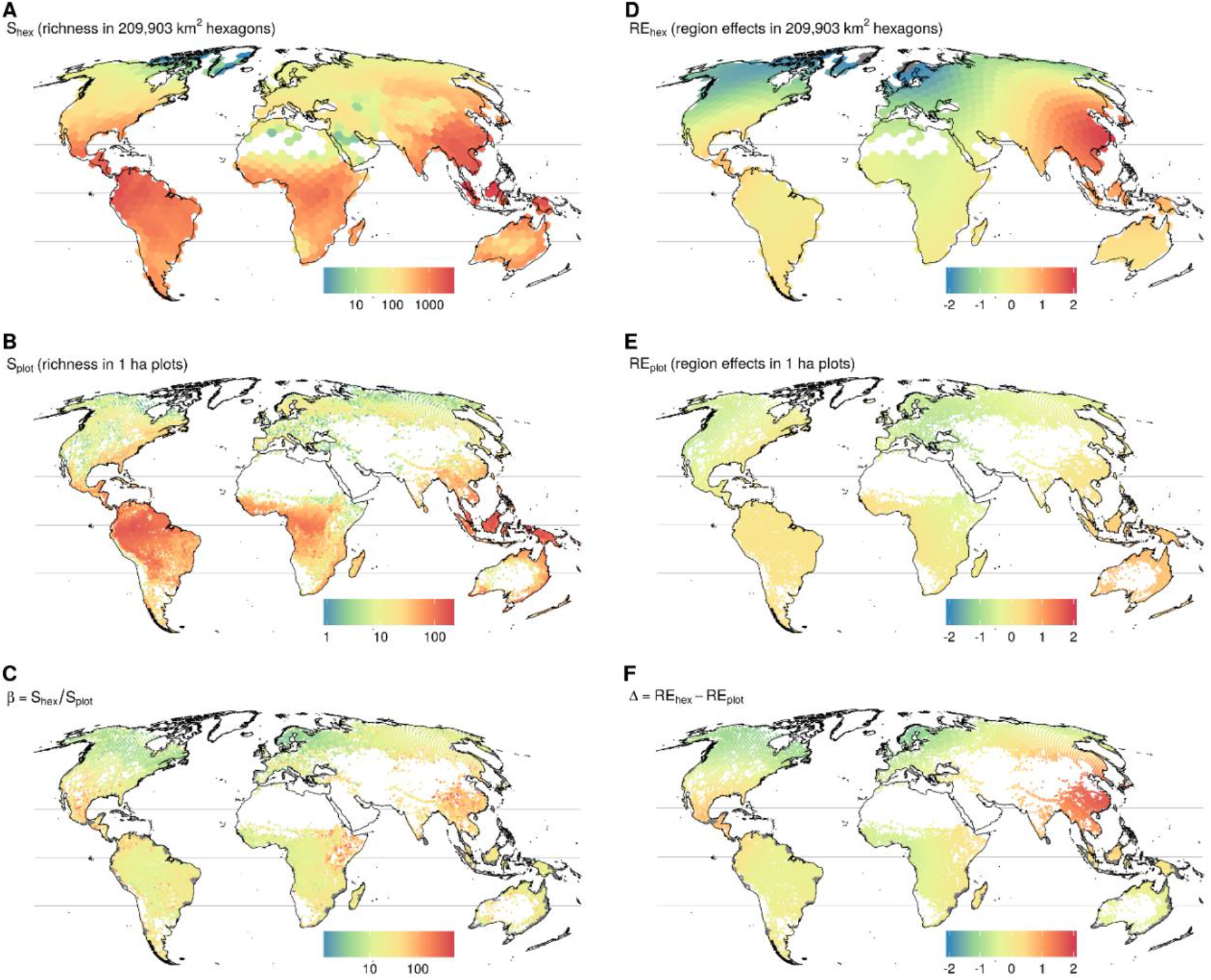
Predicted patterns of species richness, beta diversity, and continuous region effects from model SMOOTH at two spatial grains. Species richness S_hex_ is regional gamma diversity (A), S_plot_ is alpha diversity (B), and their ratio is beta diversity (C). Regional effects RE (D-E) are smooth splines representing anomaly of S (on natural log scale) from expectation based purely on environmental conditions. White mainland areas are those for which we lacked data on at least one predictor. Panels A-C use log_10_ scale.

### Grain-dependent effects of region

Although model REALM treats the regional biogeographic effects on S as discrete, while model SMOOTH treats them as continuous, both models reveal similar grain-dependence of these regional effects. At the coarse grains (i.e. in larger regions), model REALM shows that the anomaly of S that is independent of environment (and thus attributed to the effect of regions) is highest in the Indo-Malay region, followed by parts the Neotropics, Australasia, and Eastern Palaearctic (Fig. 3). Similar pattern emerges at the coarse grain from model SMOOTH, where particularly China, and Central America to some degree, are hotspots of environmentally-independent S (i.e., strong effects of biogeographic regions) (Fig. 2D). This follows the existing narrative^44,46^ where tree diversity is typically highest, and anomalous from the climate-driven expectation, in eastern Asia. However, at the smaller plot grain, a different pattern emerges in both the REALM (Fig. 3) and SMOOTH (Fig. 2E) models: the regional biogeographic effects are present, but weaker. Further, they shift away from the Indo-Malay and the Neotropical regions (REALM model) or China and Central America (SMOOTH model) at the coarse grains towards the equator, particularly to Australasia, at the plot grain (Fig. 2F, 3).

**Figure 3.**
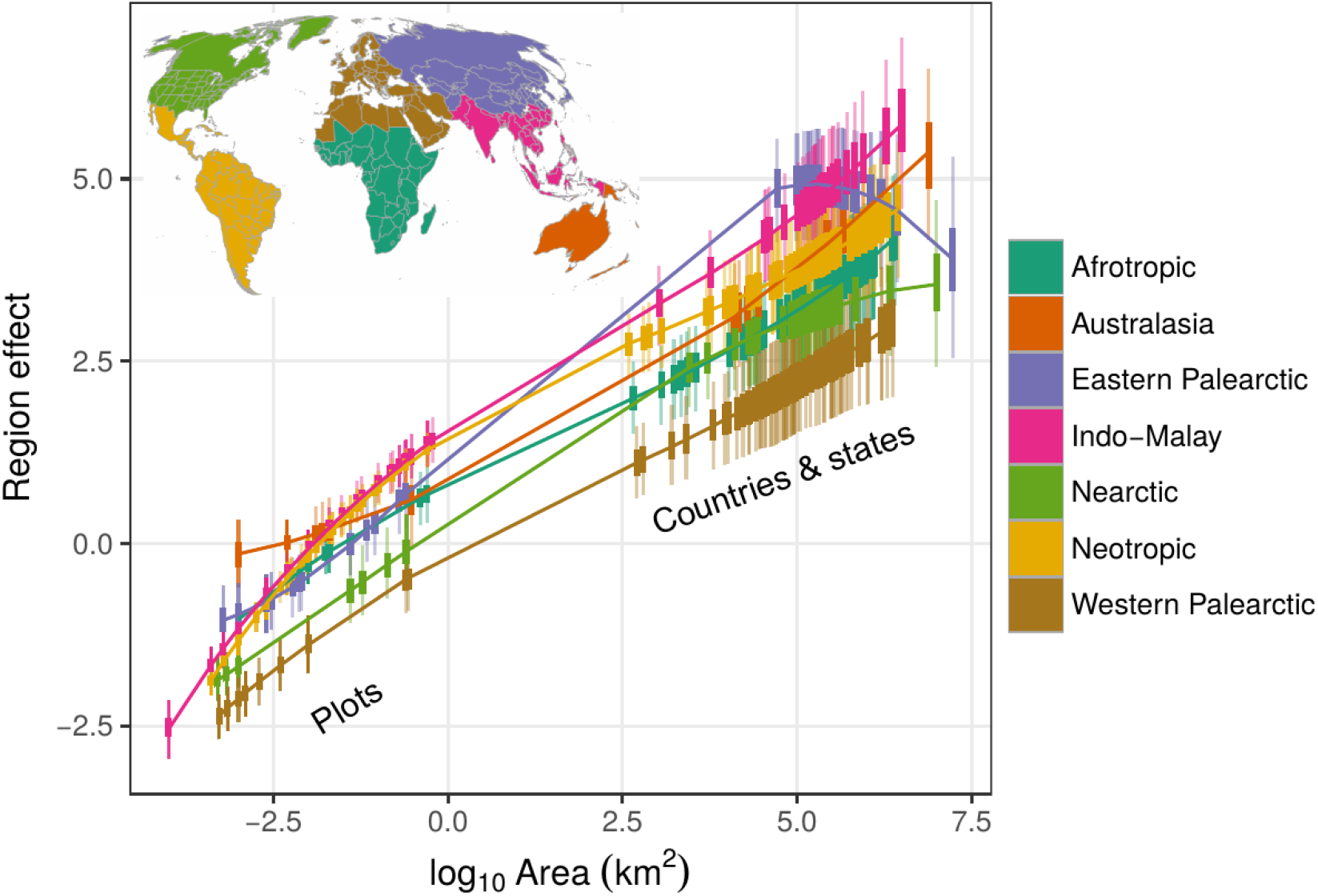
Grain-dependent effects of discrete biogeographic regions on species richness, estimated by model REALM across a continuum of areas (grains). A region effect is what remains after accounting for effects of all other predictors at a given area. Vertical bars show 2.5, 25, 50, 75 and 95.5 percentiles of posterior densities.

These results can be viewed through the logic of species-area relationship (SAR), and its link to alpha, beta, and gamma diversity^12,53^: If environmental conditions are constant, or statistically controlled for, then S depends only on area and on specific regional history. Since these interact, what emerges are region-dependent SARs (in model REALM; Fig. 3), which are equivalent to grain-dependent effects of regions (in model SMOOTH; Fig. 2). In both, what geographically varies is the environmentally-independent local (RE_plot_ in Fig. 2E) S and regional (RE_hex_, Fig. 2D) S, and their ratio (i.e. difference in log space in Fig. 3F), which directly links to the slopes of relationships in Fig. 3. We can explain this through different range dynamics in different parts of the world. Areas with high levels of environmentally-independent S at large grains, such as China and Central America, could have historically accumulated species that are spatially segregated with relatively small ranges, for example due to allopatric speciation^44^, climate refugia [as in Europe^54^], or due to dispersal barriers and/or large-scale habitat heterogeneity^44^. This would lead to increased regional richness but contribute less to local richness, leading to stronger regional effects at larger than smaller grains, as we observed.

We also found pronounced autocorrelation in the residuals of the REALM model at the country grain, but low autocorrelation at both grains in the residuals of model SMOOTH (Supplementary Fig. 5). Residual autocorrelation in S is the spatial structure that was not accounted for by environmental predictors; it can emerge as a result of dispersal barriers or particular evolutionary history in a given location or region^55,56^. The autocorrelation in REALM residuals thus indicates that the discrete biogeographical regions (Fig. 3A) fail to delineate areas with unique effects on S; these are better derived directly from the data, for example using the splines in model SMOOTH (Fig. 2D, E). As such, the smoothing not only addresses a prevalent nuisance [i.e. biased parameter estimates due to autocorrelation^57^], but can also be used to delineate the regions relevant for biodiversity more accurately than the use of á priori defined regions.

### Grain-dependent effects of environment

Generally, the signs and magnitudes of the standardized coefficients of environmental predictors (Fig. 4) at the plot grain are in line with those observed^46^. However, as far as we are aware, only Kreft & Jetz^36^ modelled richness-environment associations as grain-dependent by using the statistical interactions between an environment and area. In our analyses, several of these interactions terms were significant in both models REALM and SMOOTH (Fig. 4); this in line with refs^21–23^, but it contrasts with Kreft & Jetz^36^ who detected no interaction between area and environment at the global extent in plants. However, the latter study lacked data from local plots (i.e. had a limited range of areas). We detected the clearest grain dependence in the effect of Gross Primary Productivity (GPP, a proxy for energy input) and Tree density (Fig. 4); both effects decrease with area. The reason is that, as area increases, large parts of barren, arid, and forest-free land are included in the large countries such as Russia, Mongolia, Saudi Arabia, or Sudan, diluting the importance of the total tree density at large grains.

**Figure 4.**
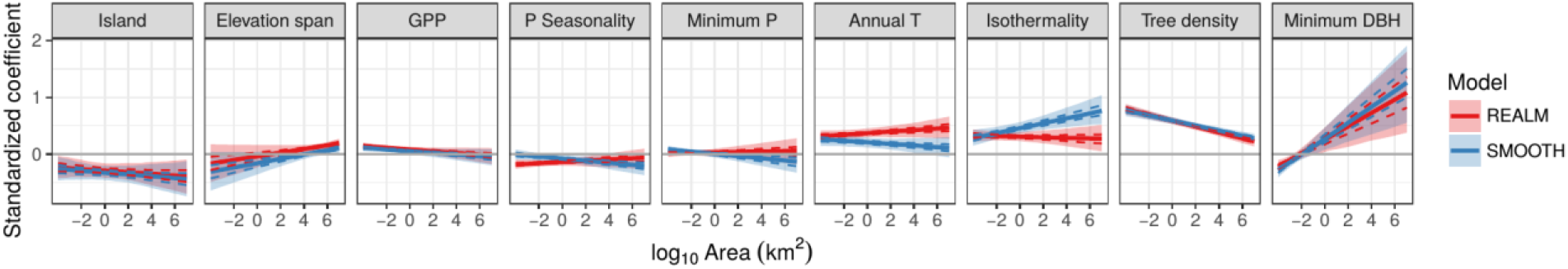
Grain-dependent standardized model coefficients for environmental predictors. The y-axis is the value of model coefficient, not S -- although the lines for tree density are declining, its effect on S is positive at all values of area. Lines and shadings are 2.5, 25, 50, 75 and 95.5 percentiles of posterior densities.

Further, we failed to detect an effect of elevation span at fine grain, but it emerged at coarse grains (Fig. 4). This is in line with other studies^21,22^, and it shows that topographic heterogeneity is most important over large areas where clear barriers (mountain ranges and deep valleys) limit colonization and promote diversification^58^. Also note the wide credible intervals (i.e. high uncertainty) around the effects of islands and most of the climate-related variables across grains (Fig. 4). A likely source of this uncertainty is the collinearity between environmental and regional predictors (see below). This prevented us from detecting grain-dependency of the effect of temperature, although we expected it based on previous studies^21,23^. Finally, we detected a consistently negative effect of islands on S, but with broad credible intervals across all grains (Fig. 4); this uncertainty is likely caused by our binomial definition of islands. We suggest that inclusion of proximity to mainland or island history (Britain or the Sunda Shelf islands used to be mainland), and inclusion of remote oceanic islands, could reduce the uncertainty.

### Regions vs environment

We used deviance partitioning^59,60^ to assess the relative importance of biogeographic regions versus environmental conditions in explaining the variation of S across grains. At the global extent, the independent effects of biogeographic realms strengthened towards coarse grain, from 5% at the plot grain to 20% for country grain in model REALM (Fig. 5A). In contrast, the variation of S explained uniquely by environmental conditions (around 14%, Fig. 5A) showed little grain dependence. However and importantly, at both grains, roughly 50% of the variation of S is explained by an overlap between biogeographic realms and environment, and it is impossible to tease these apart due to the collinearity between them. In other words, biogeographic realms also tend to be environmentally distinct (Supplementary Figs. 6–7), i.e. they are not environmentally similar replicates in different parts of the world [see also ref^46^ for similar conclusion]. The same problem prevails when the World is split into two halves and when the partitioning is done in each half separately (Fig. 5B, C). This climate-realm collinearity at the global extent weakens our ability to draw conclusions about the relative importance of contemporary environment versus historical biogeography, since by accounting for environment, we inevitably throw away a large portion of the regional signal, and vice versa. Thus, we caution interpretations of analyses such as ours and others^30,31,33,46,61^ inferring the relative magnitude biogeographic versus environmental effects merely from contemporary observational data.

**Figure 5.**
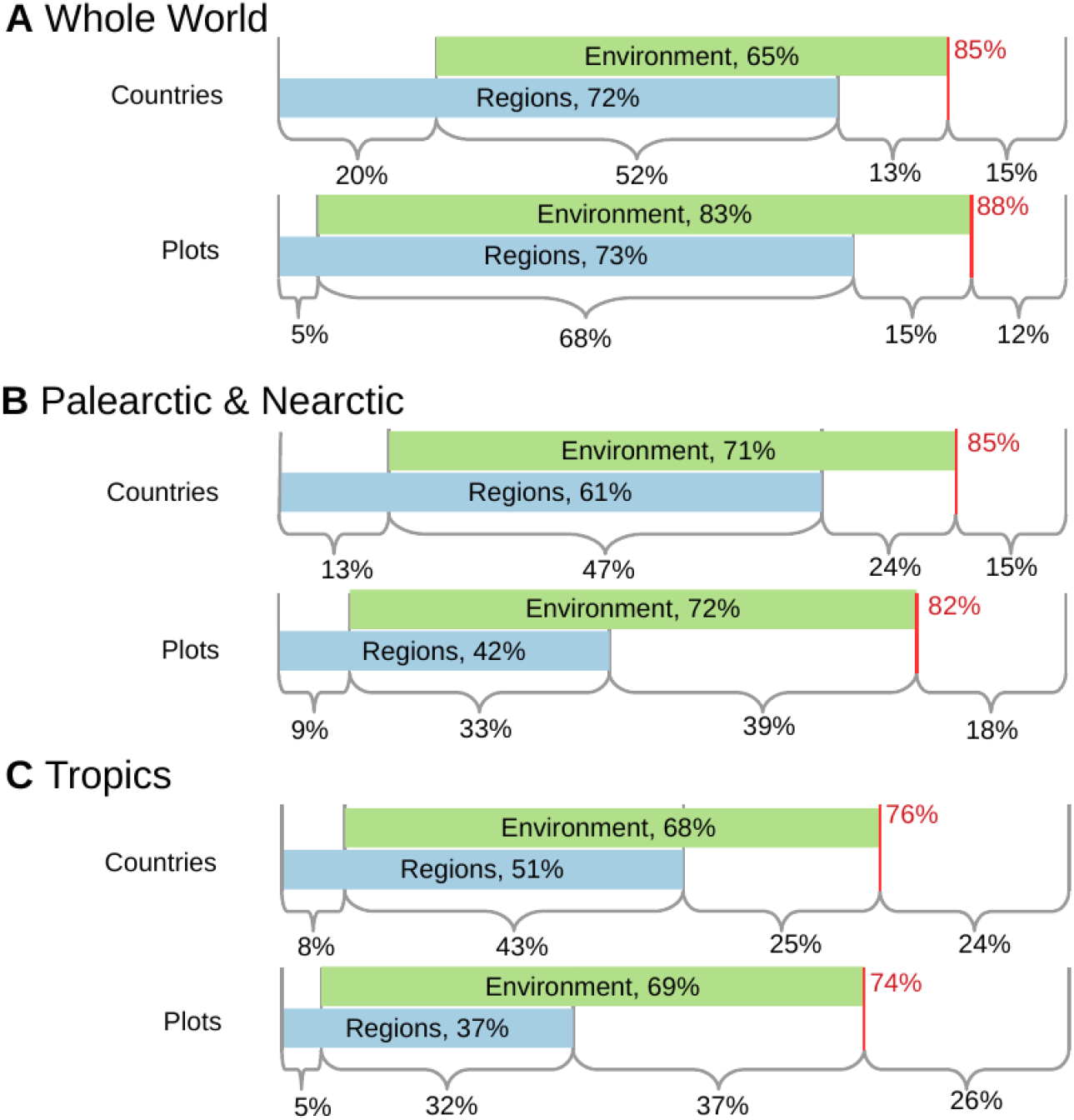
Partitioning of deviance of species richness (S) to components explained by environment vs biogeographic regions, at two spatial grains, using the REALM model. In panel A the extent is the whole World, panel B uses data from the Palearctic and Nearctic regions. Panel C uses data from the Neotropic, Afrotropic, Indo-Malay and Australasian regions. Red lines are the total deviance explained by the full model with both the environment and regions.

Given this covariation, we cannot clearly say whether environment or regional effect are more important in driving patterns of richness. We can, however, make statements about the grain dependence of both environment and region, as above. The climate-realm collinearity is likely responsible for the inflated uncertainty [as expected from ref^62^] around the effects of environmental predictors (Fig. 4) and biogeographic realms (Fig. 3), but there remains enough certainty about the effects of some predictors, such as tree density or GPP (Fig. 4), which are more orthogonal to climate and regions.

To overcome the global collinearity problem and to better answer the classical question of whether diversity is more influenced by historical or contemporary processes, we suggest the following alternative strategies: (i) analyze smaller subsets of data where environmental and regional data are less collinear, e.g. across islands^63^ or biogeographic boundaries^44,64^ with similar environment but different history, (ii) use historical data from fossil or pollen records^65^, (iii) use long-term range dynamics or other patterns reconstructed from phylogenies^66,67^, and (iv) use historical data on past environmental conditions^68^. Finally, (v) we see a promise in the emerging use of process-based and mechanistic models in macroecology^69,70^ which can predict multiple patterns, ideally at multiple grains, and as such can offer a strong tests^71^ of the relative importance of historical biogeography versus contemporary environment in generating biodiversity, irrespectively on the mutual arrangement of geography and environment.

### Implications

We have compiled a global dataset on tree species richness, and used it to integrate highly heterogeneous data in a model that contains grain-dependence as well as spatial autocorrelation, and predicts hotspots of biodiversity across grains that span 11 orders of magnitude, from local plots to the entire continents. This is an improvement of data, methods, and concepts, and importantly, we reveal a critical grain-dependence in the both regional and environmental predictors. We propose that this grain-dependence, together with the confounding collinearity between environment and geography, is the reason why studies comparing the importance of environmental versus historical biogeographic predictors of global diversity patterns have come to disparate conclusions. Studies using smaller-grained data tend to find strong influence of environment^39,40^, whereas those that use larger-grained data find strong effect historical biogeography^43,45^. We reconcile this with a grain-explicit analysis and show that smaller-grain (alpha-diversity) patterns are less strongly influenced by regional biogeography than larger-grained (gamma-diversity) patterns. Finally, we suggest that the advantages of having a formal statistical way to directly embrace grain dependence are twofold: Not only it will allow ecologists to test grain-explicit theories, but it is precisely the same grain dependence that will also allow integration of heterogeneous, messy, and haphazard data from various taxonomic groups, especially the data deficient ones. This is desperately needed in the field that has restricted its global focus to a small number of well-surveyed taxa.

## Methods

The complete data and R codes used for all analyses are available under CC-BY license in a GitHub repository at https://github.com/petrkeil/global_tree_S. Extended description of methods is in SI Text.

### Data on S at the plot grain

We compiled a global database of tree species richness in 1932 forest plots, from which we selected only plots with unique coordinates, and with data on number of individual trees, minimum diameter at breast height (DBH), and area of the plot. We included only plots that covered a contiguous area and in which all trees within the plot above the minimum DBH were determined. In case there were several plots with the exactly same geographic coordinates, we chose one plot with the largest area. If areas were the same, we chose one plot randomly. This left us with 1338 forest plots for our main analyses. Although all of these plots are in forests, the authors of the primary studies still differ in which individuals are actually determined. For instance, authors may include or exclude lianas. Thus, in the main analyses we included all plots that have the following morphological scope: “trees”, “woody species”, “trees and palms”, “trees and shrubs”, “trees and lianas”, “all living stems”. In a parallel sensitivity analysis we used a more stringent selection criteria to create a subset of the data (see below). The data are available at https://github.com/petrkeil/global_tree_S. The list of references used for data extraction is in Supplementary Information.

### Data on S at the country grain

We compiled data on tree species richness of 287 countries and other administrative units (US and Brazilian states, Chinese provinces). We downloaded the data from BONAP taxonomic data center at http://bonap.net/tdc for the United States^72^, from^73^ for the provinces of China, from Flora do Brasil 2020 at http://floradobrasil.jbrj.gov.br^74^, and from Botanic Gardens Conservation International database GlobalTreeSearch^7^ (accessed 18 Aug 2017) for the rest of the world. To download the data from GlobalTreeSearch we used Selenium software interfaced through a custom R script.

### Predictors of species richness

For each plot and each country we calculated its latitude, longitude, and area, and we extracted environmental variables (Supplementary Table 1) related to energy availability, climate seasonality, climatic limits, topographic heterogeneity, insularity, tree density, and productivity, all of which are known to predict plant and tree species richness^36,38,40,46,75^

(Supplementary Table 1). For each plot we also noted minimum DBH that was used as a criterion to include tree individuals in a study. All continuous predictors were standardized to 0 mean and unit variance prior to the statistical modelling.

### Cross-grain models

Our core approach is that ‘grain dependence’ of an effect of a predictor can be modelled as a *statistical interaction* between the predictor and area. For example, imagine a linear relationship between species richness *S* and temperature *T*, defined as *S* = *a* + *bT*. Now let us assume that the coefficient *b* also depends linearly on area (grain) *A* as *b* = *α* + *βA*; by substitution we get *S* = *a* + *α* + *βAT*, where *βAT* is the interaction term. Following this logic, we built statistical models that treat environmental and historical predictors of S as grain-dependent. Specifically, we built two models (REALM and SMOOTH) representing the same general idea of grain-dependency, but each implementing it in a somewhat different way. These models are not mutually exclusive, but are complementary approaches to the same problem.

### Model REALM

This model follows the traditional approach to assess regional effects on S, that is, variation of S that is not accounted for by environmental predictors can be accounted for by membership in pre-defined discrete geographic regions [as in^46^], a.k.a. realms. We extend this idea by assuming that the effect of biogeographic regions interacts with area (i.e. grain), i.e. there is a different species-area relationship at work in each region. Formally, species richness *S*_*i*_ in *i*th plot or country is a negative binomial random variable *S*_*i*_ ∼ *NegBin*(*μ*_*i*_, *Θ*), where

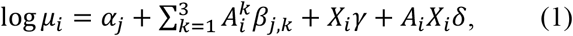

and where *α*_*j*_ are the intercepts for each *j*th region, 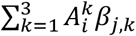 is the interaction between a third-order polynomial of area A and the *j*th region; we have chosen the third-order polynomial to ensure an ability to produce a tri-phasic effect of area^12^ in each region. *X*_*i*_*γ* is the term for area-independent effects of environmental predictors in a matrix *X*, and *A*_*i*_*X*_*i*_*δ* is the interaction term between area *A* and *X*. Parameters to be estimated are the vectors *α, β, γ, δ*, and the constant *θ*. The model can be specified in R package mgcv^76^ as gam(S ∼ REALM + poly(A,3):REALM + X + X:A, family=‘nb’), where REALM is a factor identifying the regions.

### Model SMOOTH

In this model we avoid using discrete biogeographic regions; instead, we use thin-plate spline functions (a.k.a. splines)^76^ of geographic coordinates. This allows us (i) to identify the areas of historically accumulated *S* directly from the data, and (ii) to account for spatial autocorrelation in model residuals^57^ at the same time. As above, *S*_*i*_ ∼ *NegBin*(*μ*_*i*_, *Θ*), but

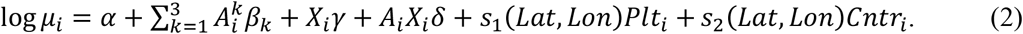

The notation is the same as in the previous model, with the exception of *α* and *β* now being constant, and with the spline functions *s*_1_ and *s*_2_ (each with 14 spline bases), and with *Plt*_*i*_ and *Cntr*_*i*_ as binary (0 or 1) variables specifying if an observation *i*is a country or a plot. In R package mgcv the model writes as gam(S ∼ s(Lat, Lon, by=Plt.or.Cntr, bs=’sos’, k=14) + poly(A, 3) + X + X:A, family=’nb’), where Plt.or.Cntr is a factor identifying if an observation is a plot or a country.

### Model fitting, inference, predictions, and sensitivity analysis

We used a combination of maximum likelihood (fast, easy to work with) and Hamiltonian Monte Carlo (slow, but handles uncertainty well) to optimize and fit the models. To compare the effects of contemporary environment vs biogeographic regions, we used partitioning of deviance. We used model SMOOTH to generate the global predictions (Fig. 2) in a set of artificially generated plots (each with an area of 1 ha) and hexagons (each with an area of 209,903 km2). We additionally tested if our results are sensitive to data sources and definition of what a ‘tree’ is. All these steps are described in detail in SI Text.

## Acknowledgements

We thank to Dylan Craven and Irena Šímová for valuable advice, and to Robert Ricklefs, Shane Blowes and two anonymous referees for critical comments that greatly improved early versions of the manuscript. We acknowledge the support of the German Centre for Integrative Biodiversity Research (iDiv) Halle-Jena-Leipzig funded by the German Research Foundation (FZT 118).

## Author contributions

P.K. formalized the ideas, collated the data, performed the analyses, and led the writing. J.M.C. proposed the initial idea, contributed to its development, discussed the results, and contributed to the writing.

